# A comprehensive assay of social motivation reveals sex-differential roles of ASC-associated genes and oxytocin

**DOI:** 10.1101/2022.05.21.492918

**Authors:** Susan E. Maloney, Simona Sarafinovska, Claire Weichselbaum, Katherine B. McCullough, Raylynn G. Swift, Yating Liu, Joseph D. Dougherty

## Abstract

Social motivation is critical to the development of healthy social functioning. Autism spectrum condition (ASC) is characterized in part by challenges with social communication and social interaction. The root of these challenges is hypothesized to be a deficit in social motivation, specifically in one or more subcomponents (e.g. social reward reward seeking or social orienting). Current social behavior assays lack the ability to quantitatively measure both social reward seeking and social orienting simultaneously. We have developed an automated socially-rewarded operant conditioning task coupled with video tracking, to quantify effort to achieve access to a social partner and concurrent social orienting behavior in mice. We established that adult wildtype mice will work for access to a social partner, that male mice exhibit greater social motivation compared to females, and there is high test-retest reliability in the task across multiple days. We then benchmarked the method with two test-case manipulations. We first tested a mouse model of Phelan-McDermid syndrome, a neurodevelopmental disorder associated with ASC. These *Shank3B* mutants failed to show social reward seeking and exhibited reduced social orienting. Next, we demonstrated that oxytocin receptor antagonism decreased social motivation in wildtype mice, consistent with its role in social reward circuitry. Intriguingly, only male mice were vulnerable to *Shank3B* mutation, while females were more vulnerable to oxytocin blockade, a double dissociation suggesting separate circuits for social motivation in male and female brain. Overall, we believe this method provides a valuable addition to the assessment of social phenotypes in rodent models of ASC and the mapping of potentially sex-specific social motivation circuits in the brain.

## Main

In social species such as humans and mice, social interactions are inherently rewarding and necessary for typical development. Social motivation, defined as the internal processes that drive these social interactions, has been conceptualized in terms of several interrelated components including social orienting (attending to social stimuli) and social reward seeking (incentive value of social interactions), with each potentially mediated by distinct brain circuits (Chevallier et al., 2012).

Rodent models are widely used to investigate the neural circuitry of social behavior. Specifically, there are well-established assays for quantifying sociability, including the three-chamber social approach test (Moy et al., 2004) and reciprocal social interaction in an open field (Ricceri et al., 2016). While valuable for the assessment of specific areas of social behavior, neither of these tasks quantify social motivation directly (i.e., the amount of work an animal will exert for social interactions). The rewarding properties of social contact can be evaluated via a social conditioned place preference (CPP) assay, in which animals learn to associate social interaction with a particular environmental context and are then given a choice between the socially-associated environment or one associated with isolation. Preference for the socially-conditioned side is thought to reflect the rewarding value of social contact (Dölen et al., 2013; Panksepp and Lahvis, 2007; Pinheiro et al., 2016). However, this assay does not measure the social orienting aspect of social motivation, and it suffers from other limitations as well: a recent review concluded that social CPP is highly variable, transiently expressed, and occurs only in specific conditions (Cann et al., 2020). Thus, while well-established procedures exist to investigate social interaction and limited aspects of social reward, methods for direct quantitative assessment of social motivation and its subcomponents are lacking. Such methods would allow more precise mapping of social motivation circuits and studies of their regulation by genes associated with human conditions.

Better understanding of social motivation circuits is important because, while challenges with social functioning are defining characteristics of Autism Spectrum Conditions (ASC), the underlying circuits remain uncertain. Recent human genetic studies have identified hundreds of ASC-associated genes (Satterstrom et al., 2020) prompting rodent studies to explore how mutations in these genes might affect conserved social circuits in the mammalian brain (Choe et al., 2022; Hu et al., 2021; Walsh et al., 2018). The Social Motivation Theory suggests that social interaction may be less motivating to individuals with ASC (Chevallier et al., 2012; Kohls et al., 2012), and that this reduced motivation may lead to poor acquisition of social skills and further social disengagement. In line with this theory, ASC children demonstrated reduced behavioral and neural responses to social rewards (Bottini, 2018; Kohls et al., 2012), and deficits in orienting to social stimuli (i.e., eye tracking) (Constantino et al., 2017; Papagiannopoulou et al., 2014). However, mechanistic studies defining the circuitry underlying each of these functions in humans are challenging, especially considering the heterogeneity of ASC: gathering enough participants with the same ASC-associated mutations, controlling for environmental confounds and genetic interactions, and interrogating neural mechanisms in a causal manner are all difficult to achieve in human research.

Thus, mouse models of ASC-associated mutations provide the opportunity to systematically investigate the role of specific genes in social motivation circuits under controlled conditions, and enable application of mechanistic circuit mapping tools not available in humans (Choe et al., 2022; Hu et al., 2021; Walsh et al., 2018). For example, mice with homozygous deletion of the *Shank3B* gene in the ASC-associated Phelan-McDermid locus show reduced social approach behavior, though it remains unclear whether this is due to a lack of social motivation (Peça et al., 2011). Interestingly, such sociability reductions have only been reported in homozygous *Shank3B* null mice, lacking two functional copies of the gene, while mice heterozygous for *Shank3B* mutation failed to show this effect despite more closely modeling the the haploinsufficiency that clinically defines Phelan-McDermid syndrome (Mitz et al., 2018). In the absence of specific behavioral assays to measure both social orienting and reward seeking aspects of social motivation, the true extent of social impairment in these mice remains unknown.

Pharmacological manipulation of neural circuitry provides a complementary approach to genetic studies. Notably, the peptide oxytocin has been shown to mediate social reward in the context of pair bonding and social preference, as well as enhancing the salience of social signals in a variety of other circuits across the brain (Froemke and Young, 2021; Shamay-Tsoory and Abu-Akel, 2016). Blocking oxytocin via administration of a receptor antagonist can reduce social reward learning (related to reward seeking behavior) in rodents (Matthews et al., 2005).e Evidence in humans suggests oxytocin may increase social orienting under specific conditions (Althaus et al., 2016, 2015). Thus, oxytocin is thought to play a critical role in the neural mechanisms underlying social motivation.

What behavioral tasks might enable more direct investigation of motivational circuits? Operant conditioning has long been employed to test motivation for many types of nonsocial stimuli such as food, sucrose, and drugs of abuse. However, only a few previous studies have attempted to use social rewards (Trezza et al., 2011). Female mice have been shown to lever-press for access to their pups (Hauser and Gandelman, 1985; Van Hemel, 1973) or for time with a sexual partner (Matthews et al., 2005). Interestingly, the latter study found that mice were also motivated to access a sex-matched stimulus animal, suggesting that social reward is sufficient to drive operant responding in the absence of parental or sexual drivers. Notably, Martin et al. developed a social operant task using lever pressing and found that BTBR inbred mice exert less effort to obtain social contact compared to C57BL/6J mice, consistent with known differences in overall sociality between these strains (Martin et al., 2014; Martin and Iceberg, 2015). A similar approach has recently been used to map social motivation circuits in wildtype mice as well (Hu et al., 2021).

Here, we present in depth a method for parallel quantitative assessment of multiple aspects of social motivation - social reward reward seeking and concurrent social orienting - by building on existing methods to couple a social operant conditioning paradigm with automated video tracking of mice. We designed a simple, inexpensive, add-on device compatible with existing operant conditioning systems and developed a significantly abbreviated conditioning protocol compared to previous studies. The training sessions are fully automated, combining automated quantification of rewarded hole-poking behavior to measure social reward seeking with simultaneous automatic video tracking of the animals’ movement to measure social orienting, by quantifying time spent in proximity to and interacting with the social partner during rewards. We first demonstrate that mice will work harder for access to a conspecific than the nonsocial condition of simply opening a door, validating the social specificity of this reward. We then assess reproducibility of the assay, noting remarkable intra-individual stability of the day-to-day measurement of social motivation. Finally, we establish construct validity and further demonstrate the sensitivity of the task with the disruption of social motivation via genetic mutation of *Shank3B* and antagonism of oxytocin receptors. This method represents an important new tool for more nuanced phenotyping of rodent social behavior and will further contribute to the mapping of social motivation circuits in the brain. Ultimately, this assay will aid in understanding the consequences of gene mutations on social circuit functions.

## Results

### Mice will work for access to a social partner

We developed an approach to quantify social reward by adapting a classic operant conditioning paradigm (**Fig. 1A)** to deliver transient visual, olfactory, and limited tactile access to a social partner in response to a conditioned behavior (i.e. nosepoking; **Fig. 1B**). We hypothesized that mice could learn to distinguish an active hole, that results in access to a social partner, from an inactive hole. We further hypothesized that the conditioned response would be greater in mice rewarded with a social partner than those that were not (**Fig. 1C**). Thus, in our first experimental cohort (Cohort 1), we examined the behavior of two groups of mice. In the experimental group, following a correct nosepoke into the active hole, a vertical door opened to reveal a novel age- and sex-matched social partner for a duration of 12 seconds. In the control group, the door opened to reveal a blank wall for 12 seconds. This group distinction allowed us to confirm that the performance of the experimental group was indeed due to the social reward.

**Figure 1.**
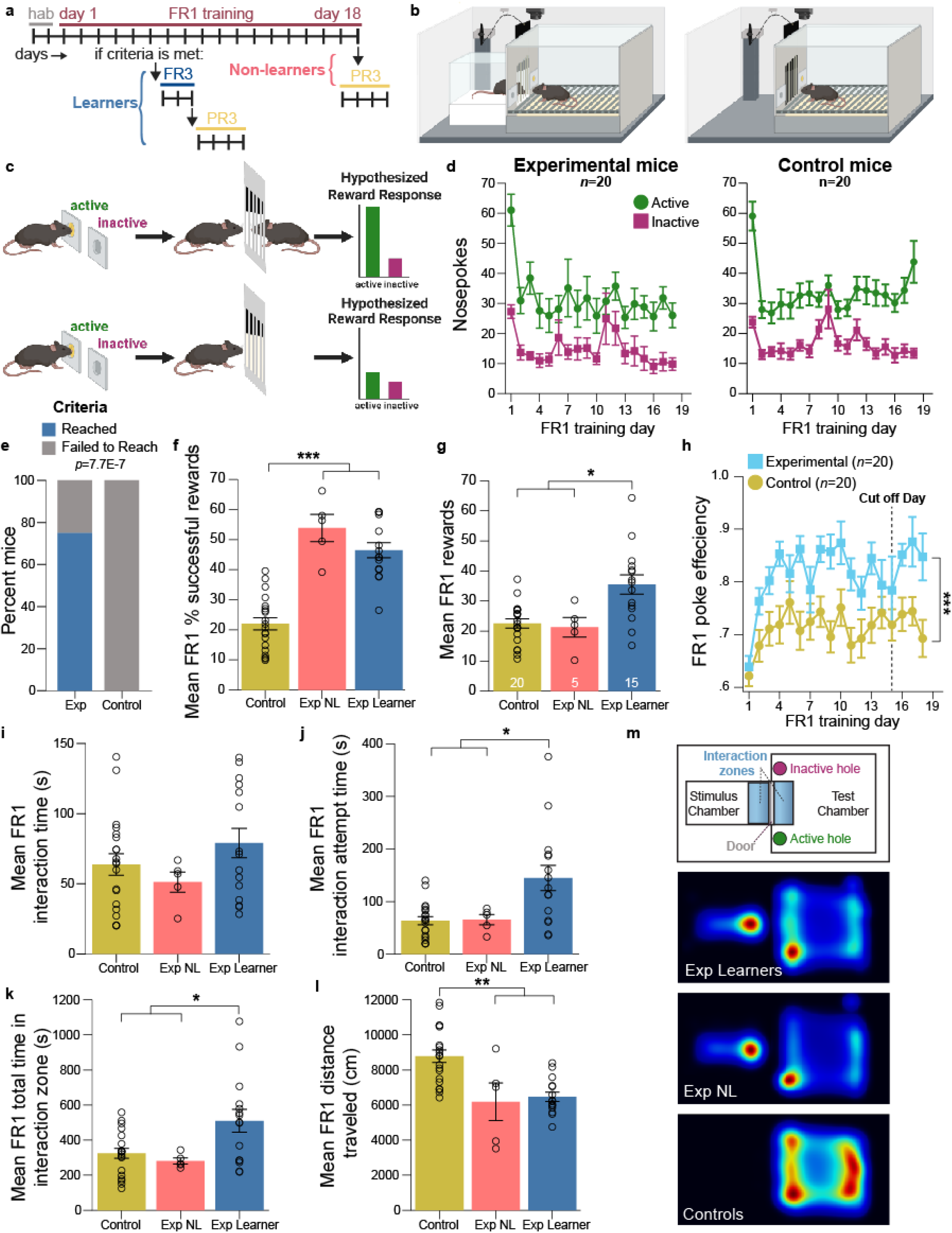
Mice exhibited both reward seeking and social orienting in the social motivation operant task. **a**. Social motivation operant task timeline schematic. **b**. Social motivation operant apparatus provides access to transient access to a social partner for the experimental condition (left panel; n=20) and access to a non-social door raising for the control condition (right panel; n=20). **c**. Schematized hypothesis that the response to reward would be greater in mice that received a social reward compared to those that did not. **d**. Nosepokes into active and inactive holes for experimental mice (left panel) and control mice (right panel) across FR1 training. **e**. Ratio of mice in experimental (15/20) and control (0/20) conditions that achieved conditioning criteria during training. **f**. Across training, the average percentage of successful rewards that included an interaction with the reward stimulus for experimental learners (Exp Learner; n=15) experimental non-learners (Exp NL; n=5) and control non-learners (Control; n=20). **g**. The average number of rewards received per day for all groups. **h**. Poke efficiency across FR1 days. Cut off day signifies the day after which no additional mice reached conditioning criteria. **i**. The duration of time spent interacting with the reward stimulus during the 12-sec reward periods averaged across FR1 days. **j**. The average time spent attempting to interact (either successfully or unsuccessfully) during FR1 training. **k**. The total time spent in the interaction zone per day averaged across FR1 training. **l**. Total distance traveled, on average, in the apparatus during FR1 training. **m**. Schematic of apparatus (top panel) and heatmaps of animals’ body positions in the apparatus across FR1 training (lower three panels). Warmer colors represent greater time spent in that position. **d**,**f-l**. Grouped data are presented as means ± SEM with individual data points as open circles. Statistical significance, ***p<.001, **p<.01, *p<.05.

It is important here to operationally define the components of social motivation assessed in this task. As defined in Chevallier et al., social reward seeking is the behavioral manifestation of the incentive value of a social reward (Chevallier et al., 2012). Thus, we operationally defined reward seeking behaviors in our task as reaching conditioning criteria, number of rewards achieved, poke efficiency (rewards/active nosepokes), percentage of successful rewards, poke accuracy (active/total nosepokes), and breakpoint (all defined in **Table 1**). Social orienting is the prioritization of attention to social signals (Chevallier et al., 2012), operationally defined here as time engaged in a social interaction with the stimulus mouse, social interaction attempts, and total time in and entries into the interaction zone (also defined in **Table 1**).

**Table 1.** Glossary of outcomes.

| Social Motivation Component | Outcome | Definition |
| --- | --- | --- |
| Seeking/liking | Conditioning criteria reached | Animal showed conditioning to the reward stimulus (criteria include at least 40 active nose pokes, 75% poke accuracy, and 65% successful rewards during a single session) |
|  | Reward | The door opening in response to the required number of active nose pokes, presenting access to a social partner and allowing for a social interaction to occur. |
|  | Poke efficiency | Rewards per active nose pokes, where a 1.0 indicates one reward was received per one active nose poke (rewards/active nose pokes). |
|  | % successful rewards | Percent of rewards where a social interaction occurred (or an interaction with the door for Cohort 1 control mice). |
|  | Poke accuracy | Percent of active nose pokes (active nose pokes/total nose pokes) |
|  | Breakpoint | Maximum nose pokes performed to elicit a reward during progressive ratio 3 testing. |
| Social orienting | Interaction | Duration of time the test and stimulus mice are simultaneously at the open door (in their respective interaction zones) during a reward |
|  | Interaction attempts | Duration of time test mouse is in the interaction zone during a reward when the door is open door, which can be both successful (an interaction occurs) or unsuccessful (the stimulus mouse is not at the door and no interaction occurs). |
|  | Total time in interaction zone | Total duration of time the test mouse is in the interaction zone during a 1-hr session, including time during and not during rewards, regardless of the stimulus mouse location. |
|  | Interaction zone entries | Entries into the interaction zone at the door. |

To determine the point at which animals were successfully conditioned to nosepoke for the reward, we defined conditioning criteria of at least 40 active nosepokes, 75% poke accuracy, and 65% successful rewards during a single session. We included the criterion of % successful rewards to ensure the behavior of the mice was driven by engagement with the reward, i.e. interaction with a social partner or the open door. All mice in both groups learned to distinguish the active from inactive holes (**Fig. 1D**) within 15 days of training on a fixed ratio schedule of reinforcement in which one active nosepoke was required to receive one reward (FR1). However, while the majority of experimental mice reached conditioning criteria, none of the control mice did (**Fig. 1E**). The control mice had a lower percentage of successful rewards on average per session compared to all experimental mice (**Fig. 1F**). In addition, the control mice received significantly fewer rewards on average per session than the subset of experimental mice who successfully conditioned (learners), performing comparably to the subset of experimental mice who did not condition (non-learners; **Fig. 1G; Table 2**). Finally, although comparable numbers of active nosepokes were achieved by experimental and control mice (**Fig. 1D**), the poke efficiency was significantly diminished among the control mice (**Fig. 1H**). This indicates that, in contrast to the experimental mice, the control mice continued to poke in the active hole even once a reward had been achieved, instead of interacting with the reward stimulus (i.e. the open door). Thus, we can conclude that the performance of the experimental mice was specifically driven by the social reward.

**Table 2.**
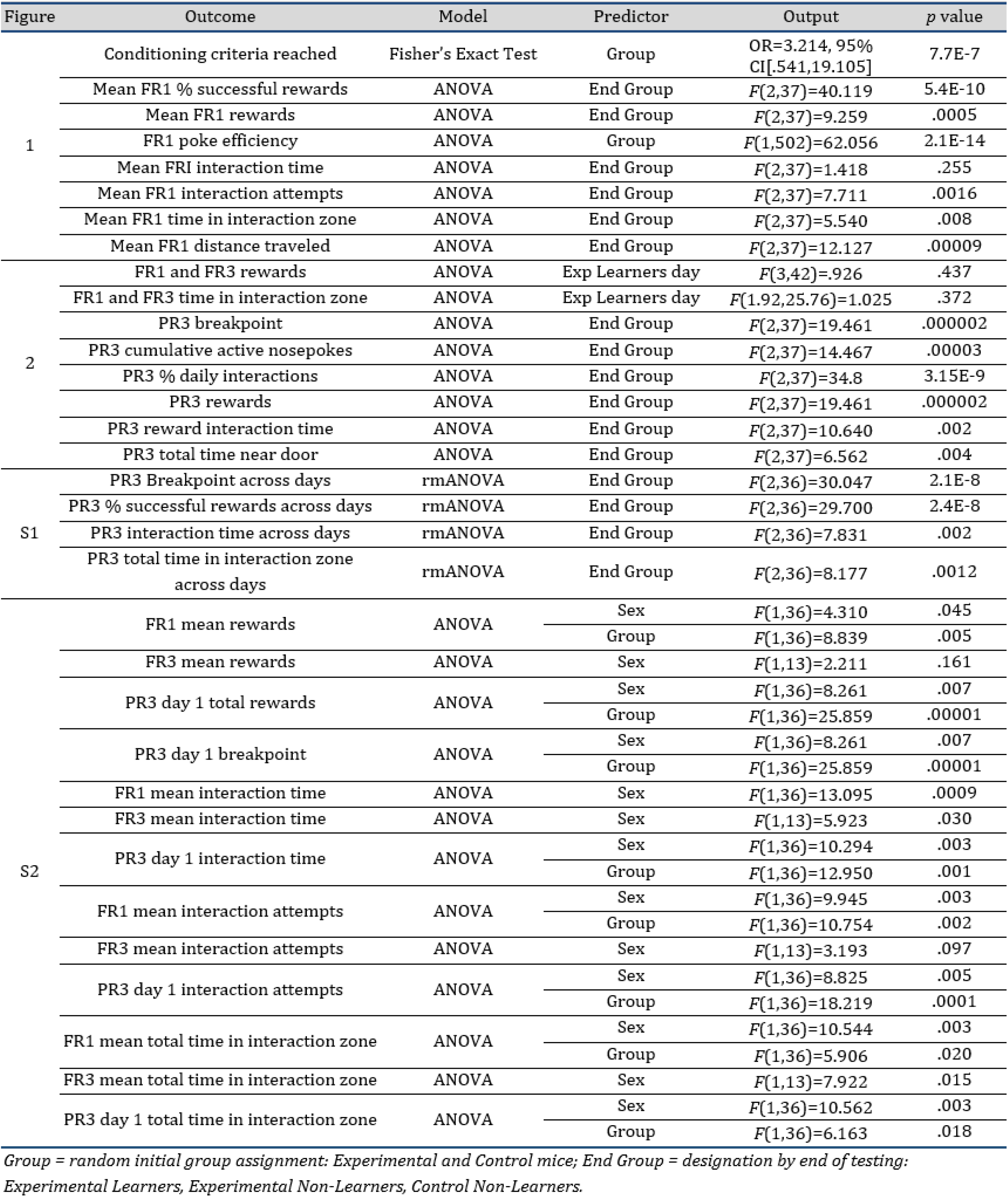
Statistical analysis results for C57BL/6J Cohort 1 experiments.

### Mice exhibited social orienting during access to a social partner

We next evaluated social orienting in our operant task. We did not observe differences in interaction time between groups (**Fig. 1I**), which is defined here as time both the test and stimulus mice are at the open door during a reward (experimental group) or time at the open door (control group). This may inflate interaction time for the control mice, however, as they don’t depend on the behavior of a second animal as in the experimental group (i.e. the stimulus mouse). Therefore, a more adequate comparison to the control group time at the open door may be made with interaction attempt time for the experimental group, or the time at the door during a reward regardless of whether the stimulus mouse is present. Here we see that experimental learners exhibited greater time attempting an interaction compared to both experimental non-learners and controls (**Fig. 1J**). In fact, experimental learners spent significantly more time in the interaction zone regardless of a concurrent reward across the entire session compared to both control and experimental non-learner mice (**Fig. 1K**). Conversely, the control mice exhibited a greater level of activity in the test chamber during the task compared to both learner and non-learner experimental groups (**Fig. 1L,M**). Together these data indicated that the experimental learners exhibited social orienting, spending time in close proximity to the social reward, while the control mice exhibited much more general exploration of the test chamber.

### Mice are highly motivated to obtain social rewards under effortful conditions

Since the majority of experimental mice were successfully conditioned to nosepoke for a social reward during training under an FR1 reinforcement schedule, we next increased the effort required to obtain this reward. In the session following their achievement of conditioning criteria, learners were moved up to a fixed ratio 3 (FR3) schedule of reinforcement, requiring three active nosepokes to achieve one reward. During FR3 training, the mice continued to show a high poke accuracy (**Fig. 2A**) and achieved comparable numbers of rewards per day as during FR1, despite the required effort increasing threefold (**Fig. 2B**). In addition, the mice spent a similar amount of time in the interaction zone on each day of FR3 compared to the average time per day during FR1 (**Fig. 2C**). Thus, when the task became more effortful, the mice continued to display high levels of social reward seeking and social orienting.

**Figure 2.**
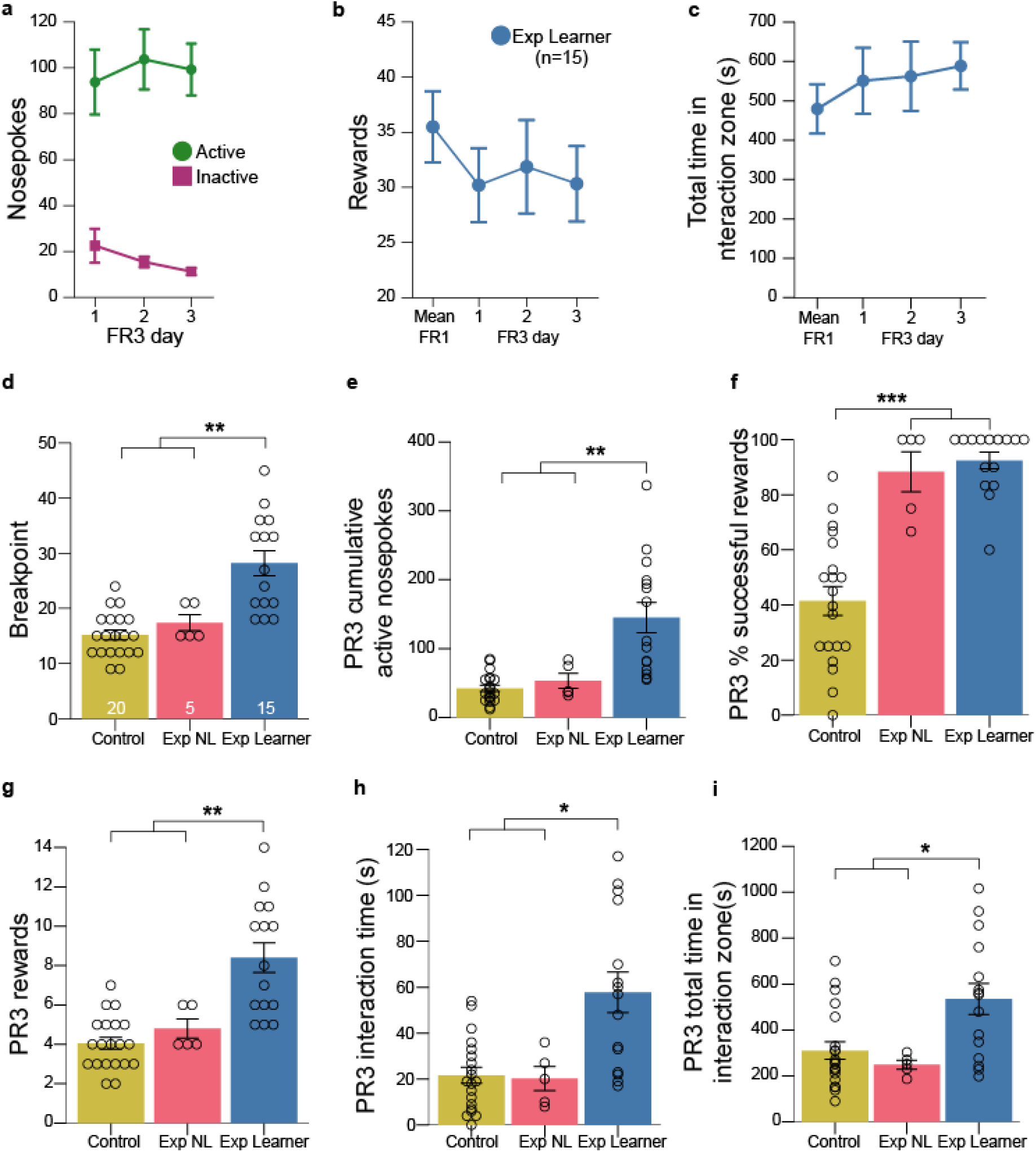
Mice were highly motivated to obtain a social reward. **a**. Nosepokes into active and inactive holes made by experimental learners (n=15) across FR3 training. **b**. Rewards achieved by experimental learners (Exp Learners) averaged across FR1 days and on each FR3 day. **c**. The total time per day in the interaction zone for Exp Learners averaged across FR1 days and on each FR3 day. **d**. Breakpoint achieved during day 1 of PR3 for Exp Learners (n=15), experimental non-learners (Exp NL; n=5) and control (Control; n=20) non-learners. **e**. Cumulative number of active nosepokes across groups on day 1 of PR3. **f**. The percent of successful rewards on PR3 day 1 across groups. **g**. The number of rewards achieved on day 1 of PR3 for all groups. **h**. The duration of interaction time on PR3 day 1 for all groups. **i**. Total time in the interaction zone across the entire PR3 day 1 session for all groups. For all panels, the grouped data are presented as means ± SEM with individual data points as open circles. Statistical significance, ***p<.001, **p<.01, *p<.05.

Finally, after three sessions at FR3, we further increased the effort required to obtain social rewards by moving to a progressive ratio 3 schedule of reinforcement (PR3), in which the number of nosepokes required increases by 3 pokes after each reward (i.e., 3,6,9,12…). By quantifying the breakpoint, or number of nosepokes at which the animal stops pursuing further rewards, we directly measured how hard the animal was willing to work for access to a social partner. Experimental learner mice achieved a significantly greater breakpoint compared to control mice and experimental non-learner mice (**Fig. 2D**), reaching up to 350 active nosepokes within a 1 hour session (**Fig. 2E**), suggesting a strong motivation to seek a social reward. Furthermore, the experimental mice, independent of learner status, achieved 88% successful rewards (learners, 92.4%; non-learners, 88.3%) whereas the control mice only achieved 41.4% successful rewards (**Fig. 2F**). Due to the much higher number of rewards received by the experimental learner group relative to the other two groups, the interaction time with the stimulus (social partner or door) was significantly greater in the experimental learner group compared to both experimental non-learners and controls (**Fig. 2G,H**). These data suggest that mice are more highly motivated to interact with a social stimulus in this paradigm compared to a control stimulus, regardless of learner status, although there appears to be a social motivation dichotomy within the mice driven at least in part by social orienting differences. As observed in FR1 training, the experimental non-learners and the controls spent significantly less total time in the interaction zone during the PR3 testing compared to experimental learners (**Fig. 2I**).

### Social motivation measurements show high reliability across days

We next explored the measurement reliability of the task within individuals across time. To do this, we continued PR3 testing for an additional three days. Breakpoint remained consistent across sessions for all groups (**Fig. S1A**) and was highly correlated within an individual (**Fig. S1B**). Further, measures of social orienting remained consistent: each group showed similar percentages of successful rewards across all four PR3 days, with the two experimental groups both significantly higher than controls (**Fig. S1C**). Similar consistency was observed for interaction time across PR3 days (**Fig. S1D**), as well as for total time in the interaction zone (**Fig. S1E**).

### Male mice exhibit more robust social reward seeking and social orienting than females

Understanding innate sex differences in social behavior can help us decipher the origins of sex biases observed in conditions like ASC (Loomes et al., 2017). Thus, we examined the influence of sex on social motivation in our data, collapsing across learner and non-learner experimental groups to enhance statistical power. Males achieved significantly more rewards on average during FR1 training and PR3 testing compared to females, though this difference was non-significant for FR3 training (**Fig. S2A-C**). Correspondingly, males reached a higher PR3 breakpoint (**Fig. S2D**). Males also showed greater interaction time, interaction attempt time, and total time in the interaction zone (**Fig. S2E-M**). In contrast, no effect of sex was observed for poke accuracy, total nosepokes, or distance traveled. In addition, learner status was not associated with sex, nor was the average number of days to reach conditioning criteria different between sexes (Females *M*=5.0 days, *SD*=1.9; Males *M*=6.4 days, *SD*=4.5). Together, these data indicate that learning and general activity levels in this task were comparable between the sexes, yet reward seeking and social orienting behaviors were greater in male mice compared to females.

### *Shank3B* mutation disrupts social motivation

*Shank3* haploinsufficiency is a highly penetrant monogenic cause of ASC (Soorya et al., 2013). Mouse models of SHANK3 loss have been well-validated, with KO mice demonstrating self-injurious repetitive grooming and deficits in social approach, social novelty, and communication (Guo et al., 2019; Peça et al., 2011; Wang et al., 2017). To provide new insights into social motivation in an ASC-associated model, we assessed performance of both male and female *Shank3B* homozygous (KO) and heterozygous (Het) mutants alongside wildtype littermates (WTs) in our social operant task (**Fig. 3A**).

**Figure 3.**
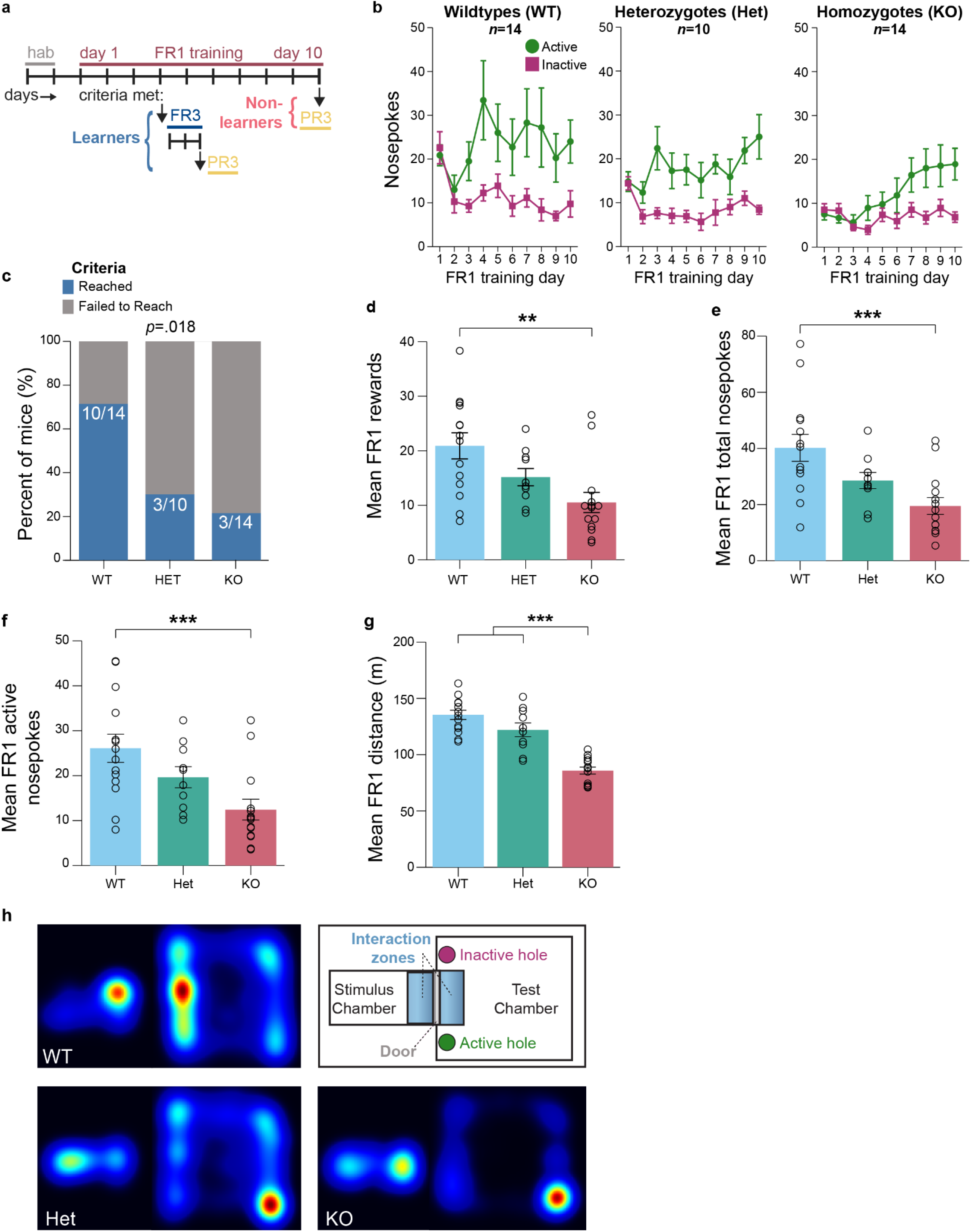
*Shank3B* knockout (KO) and heterozygous (Het) mice exhibited reduced social reward seeking and exploratory behavior relative to wildtype (WT) controls, **a**. *Shank3B* social motivation experiment timeline schematic, **b**. Nosepokes into active and inactive holes for WT (left panel, n=14), *Shank3B* Hets (middle panel, n=10), and KO mice (right panel, n=14) across FR1 training **c**. Percent of WT (10/14), Het (3/10), and KO (3/14) mice that achieved conditioning criteria during FR1 training, **d**. The number of rewards, averaged across FR1 days, attained by WT, Hets, and KOs. **e**. Mean number of total nose-pokes across FR1 days by WT, Hets, and KOs. **f**. Mean number of active nose-pokes across FR1 days by WT, Hets, and KOs. **g**. Mean distance traveled by WT, Hets, and KOs across FR1 training, h. Apparatus schematic (upper right corner) and representative heatmap of the animal’s position in the apparatus, with a single animal during a single FR1 session represented in WT (left panel) Het (middle), and KO (right) Warmer colors represent greater time spent in session represented in WT (left panel) Het (middle), and KO (right) Warmer colors represent greater time spent in that position, **d-g**. Grouped data are presented as means ± SEM with individual data points as open circles. Statistical significance, ***p<.001, **p<.01, *p<.05.

First, we sought to replicate previous findings of social approach deficits among *Shank3B* KO mutants (Guo et al., 2019; Peça et al., 2011; Wang et al., 2017). Our operant conditioning protocol includes a habituation phase prior to FR1 training, during which the test animal is allowed uninterrupted access to a novel social stimulus mouse, as the door remains raised and the nosepoke holes are inaccessible. Thus, the animal’s entries into and total time in the interaction zone provide measures of social approach behavior. As expected, KO mice made fewer entries into the interaction zone compared to WT mice (**Fig S3A, Table 3**). KO mice also traveled a shorter distance exploring the chamber than WT or Het mice, staying mainly in the corners of the test chamber furthest from the door (**Fig S3B**). A sex effect was observed for time spent in the interaction zone, with male KO mice spending significantly less time inside it compared to WT and Het males (**Fig S3B**), a pattern not observed in females. Thus, in the context of freely available access to a social partner, we observed reduced social approach among male *Shank3B* KO mice but not Het mice, similar to established phenotypes (Guo et al., 2019; Peça et al., 2011; Wang et al., 2017).

**Table 3.** Statistical analysis results for *Shank3B* Cohort 2 experiments.

| Figure | Outcome | Model | Predictor | Output | *p* value |
| --- | --- | --- | --- | --- | --- |
| S3 | Habituation entries into interaction zone | ANOVA | Genotype | $F(2,32)=11.011$ | .0002 |
| | | | Sex | $F(1,21)=12.072$ | .001 |
| | Habituation distance traveled | ANOVA | Genotype | $F(2,35)=26.966$ | $8.18E^{-8}$ |
| | Habituation time in interaction zone | ANOVA | Genotype*Sex | $F(2,32)=4.570$ | .018 |
| 3 | Criteria Reached | Pearson Chi-Square | Genotype | $X^2(2)=7.995$ | .018 |
| | Mean FR1 rewards | ANOVA | Genotype | $F(2,35)=6.872$ | .003 |
| | Mean FR1 total nose pokes | ANOVA | Genotype | $F(2,35)=8.044$ | .001 |
| | Mean FR1 active nose pokes | ANOVA | Genotype | $F(2,35)=7.039$ | .003 |
| | Mean FR1 distance traveled | ANOVA | Genotype | $F(2,35)=38.920$ | $1.27E^{-9}$ |
| 4 | Criteria Reached | Pearson Chi-Square | Genotype, Male | $X^2(2)=16.354$ | .0003 |
| | Mean FR1 interaction time | ANOVA | Genotype | $F(2,32)=3.840$ | .032 |
| | | | Genotype*Sex | $F(2,32)=6.334$ | .005 |
| | Mean FR1 interaction attempts | ANOVA | Genotype | $F(2,32)=5.351$ | .010 |
| | | | Genotype*Sex | $F(2,32)=4.292$ | .022 |
| | Mean FR1 total time in interaction zone | ANOVA | Genotype*Sex | $F(2,32)=5.849$ | .007 |
| | Mean FR1 investigation zone entries | ANOVA | Genotype | $F(2,32)=23.418$ | $5.43E^{-7}$ |
| | | | Genotype*Sex | $F(2,32)=5.449$ | .009 |
| | PR3 breakpoint | ANOVA | Genotype | $F(2,32)=5.232$ | .011 |
| | | | Sex*Genotype | $F(2,32)=5.309$ | .010 |
| | PR3 interaction attempts | ANOVA | Sex Genotype | $F(2,32)=6.134$ | .006 |
| | | | Genotype | $F(2,32)=3.170$ | .055 |
| | PR3 investigation zone entries | ANOVA | Genotype | $F(2,35)=9.248$ | .001 |

Next, we probed social motivation in *Shank3B* mice. Across FR1 training, all mice eventually learned to distinguish active from inactive nosepokes (**Fig 3B**). However, more WT mice reached conditioning criteria compared to Het and KO mice, an effect driven by male mice (**Fig 3C, 4A**). Additionally, WT mice obtained significantly more rewards on average across FR1 training than KOs (**Fig 3D**) and exhibited greater overall nosepoking than KOs, with correspondingly more active nosepokes (**Fig 3E and F**). This was likely driven by KOs’ propensity to remain inactive in the apparatus corner, as shown by their shorter distance traveled (**Fig 3G and H**). For all of these outcomes, the Het mice showed an intermediate phenotype between WT and KO, but the differences did not reach statistical significance. Together, these data demonstrate reduced social reward seeking among mice with complete loss of *Shank3B*.

Examination of social orienting behavior revealed that WTs spent significantly more time interacting during rewards than KOs, driven by WT males interacting longer than KO males likely due to the greater number of awards attained by WT males (**Fig 4B**). Relative to WT males, male Hets and KOs spent less time attempting interactions (**Fig 4C**) and less total time in the interaction zone (**Fig 4D**), with no effect observed in the females. Furthermore, entries into the interaction zone were also significantly greater for WTs than Het and KO mice (**Fig 4E**), with WT females entering the interaction zone significantly more than KO females, and WT males more than both Het and KO males (**Fig 4F**). Thus, both male Het and KO mutants exhibited reduced social orienting.

**Figure 4.**
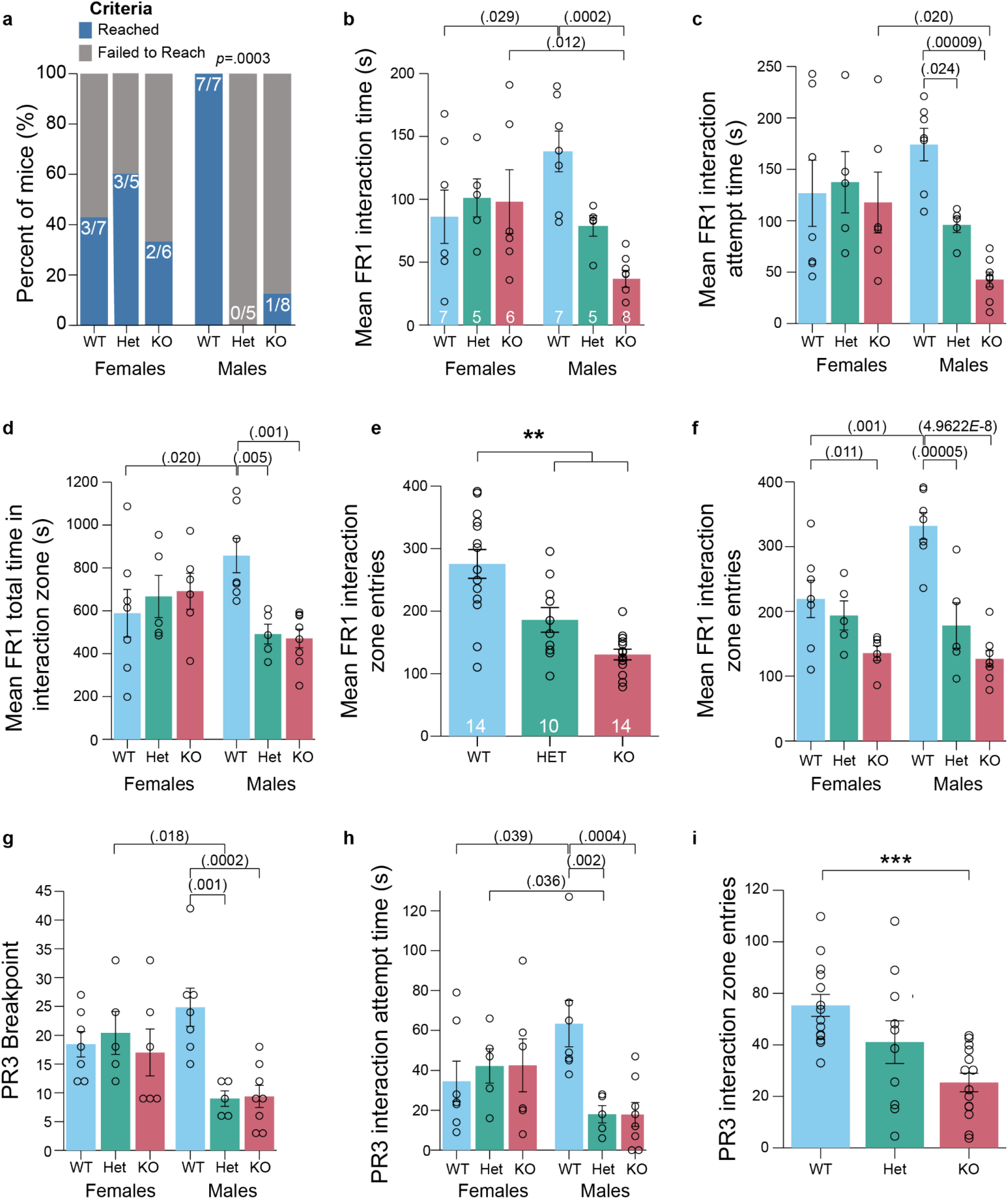
Male *Shank3B* knockout (KO) and heterozygous (Het) mice exhibited reduced social orienting and reduced social motivation relative to wildtype (WT) controls. **a**. Split by sex, percent of female WT (3/7), Het (3/5), and KO (2/6) mice, and male WT (7/7), Het (0/5), and KO (1/8) mice that achieved conditioning criteria during training. **b**. Mean duration of interaction time across FR1 training, split by sex, for WT (n=14, female=7), *Shank3B* Het (n=10, female=5), and KO (n=14, female=6). **c**. Average time attempting interactions in FR1 training. **d**. Total time in the interaction zone in FR1 training. **e-f**. Mean number of interaction zone entries across FR1 days, represented by (**e**) genotype and by (**f**) genotype by sex. **g**. Breakpoint achieved during PR3 testing. **h**. Interaction attempt time during PR3 testing. **i**. Number of entries into the investigation zone during PR3 testing. **a-h**. Grouped data are presented as means ± SEM with individual data points as open circles. Statistical significance, ***p<.001, **p<.01, *p<.05.

Finally, when reward value was directly measured in PR3 testing, male WTs achieved higher breakpoints compared to both male Hets and KOs (**Fig 4G**). WT males also spent more time attempting interactions than Het and KO males in PR3 testing (**Fig 4H**). WTs entered the interaction zone significantly more throughout PR3 testing, regardless of sex (**Fig 4I**). Overall, these findings suggest that reward seeking and social orienting are both reduced in *Shank3B* Het and KO mice compared to WTs, particularly among males. The novel observation of reduced social motivation in *Shank3B* Hets, which mimic the patient haploinsufficient phenotype, suggests a greater sensitivity of our operant task relative to existing social behavior assays.

### Oxytocin receptor blockade reduced social reward seeking but not social orienting behavior

Finally, we tested the sensitivity of this task in the context of pharmacological manipulation of neural circuits involved in social interactions. Specifically, we tested whether the oxytocin system participates in the social reward seeking and/or the social orienting aspects of social motivation by assessing these behaviors in the presence of an oxytocin receptor (OTR) blockade. Wildtype mice were administered either an OTR antagonist (OTRA) or vehicle via intracerebroventricular infusion daily prior to social operant sessions (**Fig. 5A**). During FR1 training, both infusion groups learned to distinguish between the active and inactive holes (**Fig. 5B**), with no difference in the average poke accuracy (**Fig. 5C; Table 4**). However, we observed a significant reduction in average daily total nosepokes among OTRA-infused mice compared to vehicle-infused controls (**Fig. 5D**). In addition, 61% of the vehicle-infused mice reached conditioning criteria while only 7% of OTRA mice did (**Fig. 5E**). Across FR1 training, OTRA-infused mice achieved significantly fewer daily rewards on average compared to vehicle-infused mice (**Fig. 5F**). Despite receiving fewer rewards, the OTRA-infused mice and vehicle-infused mice exhibited comparable % successful rewards (**Fig. 5G**). Finally, OTRA infusion did not alter poke efficiency (**Fig. 5H**) despite reducing the average daily total of active nosepokes (**Fig. 5I**). These data indicate that blocking oxytocin receptor activity reduced social reward seeking.

**Table 4.**
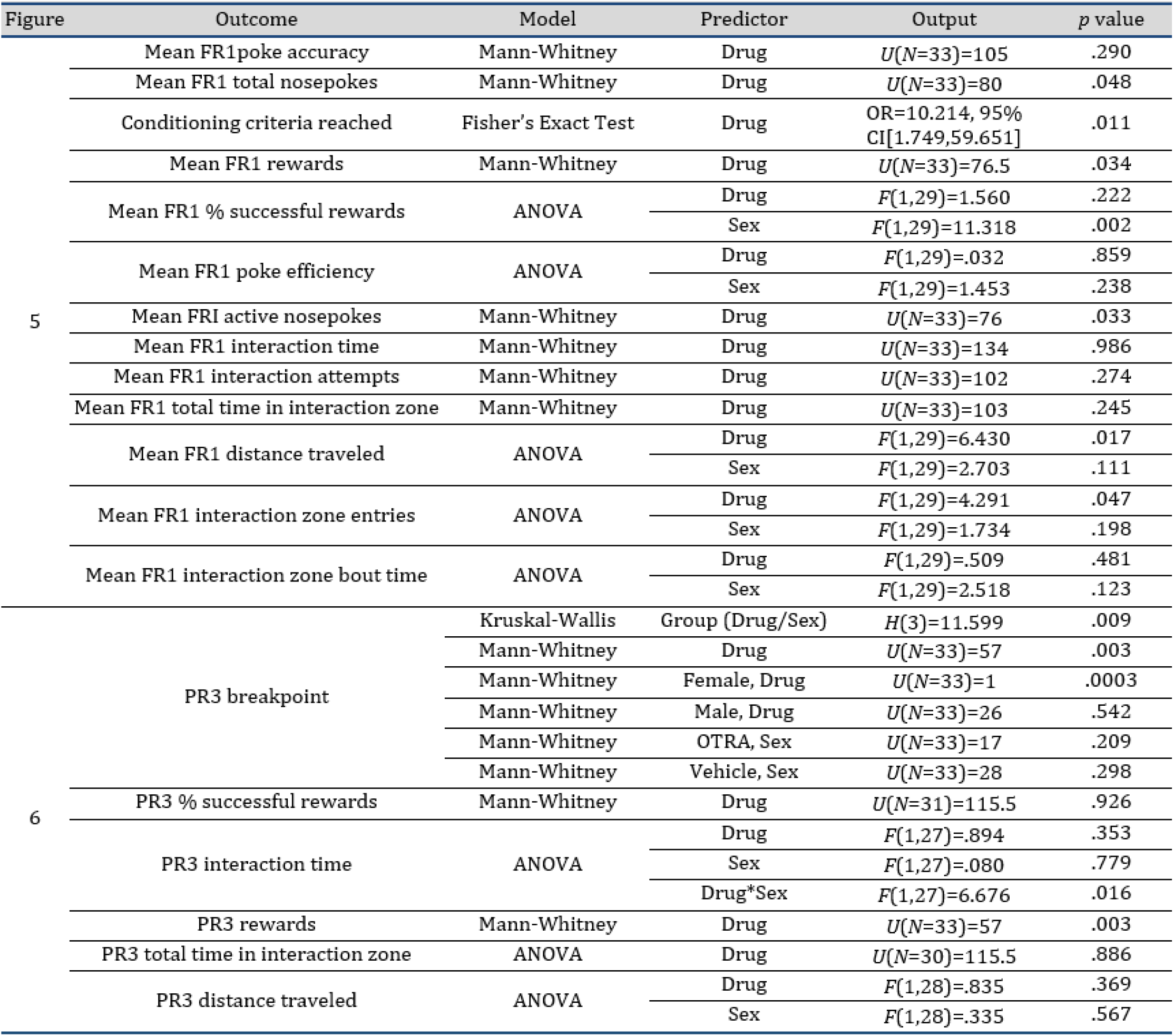
Statistical analysis results for OTRA Cohort 3 experiments.

**Figure 5.**
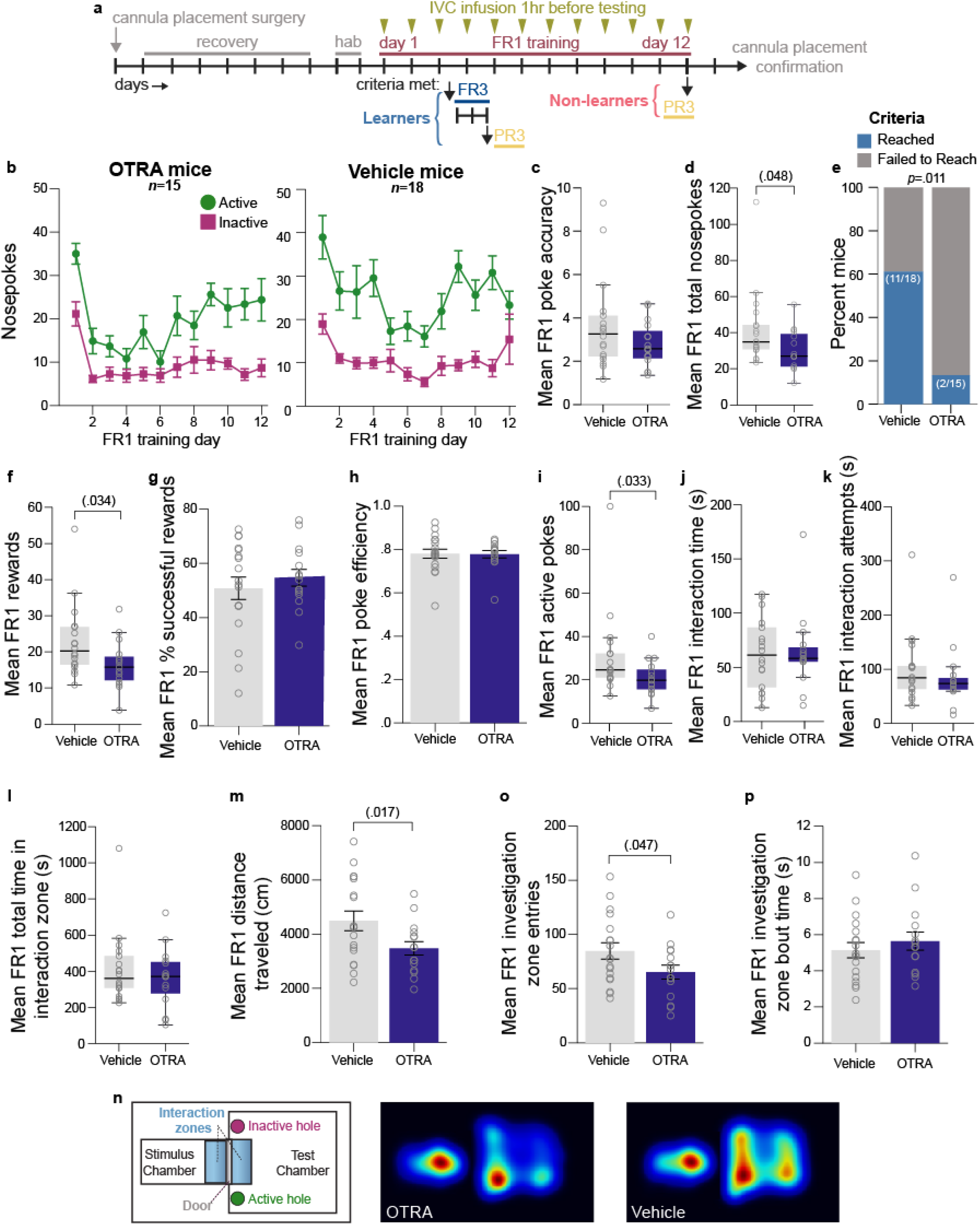
Blockade of OTRs reduced social reward seeking behavior. **a**. Experimental timeline for cannula placement surgery, social motivation operant assay with daily OTRA or vehicle infusions, followed by cannula placement confirmation via IVC dye infusion. **b**. Nosepokes into active and inactive holes for OTRA-infused mice (left panel; n=15) and vehicle-infused mice (right panel) across FR1 training. **c**. Average poke accuracy for FR1 training. **d**. Average total nosepokes for FR1 training. **e**. Ratio of OTRA (2/15) and vehicle (11/18) mice that achieved conditioning criteria. **f**. The number of average rewards received across FR1 days by both infusion groups. **g**. Across FR1 days, the average percent of successful rewards across groups. **h**. Poke efficiency across FR1 training. **i**. Average number of active nose pokes exhibited during FR1 training. **j**. Average interaction time during FR1 training. **k**. Interaction attempt time during FR1 training. **l**. Average total time in the interaction zone during FR1 training. **m**. Average distance traveled across FR1 days. **n**. Apparatus schematic (left-most panel) and heatmaps of animals’ body positions in the apparatus across the first three FR1 days. Warmer colors represent greater time spent in that position. **o**. Investigation zone entries averaged across FR1 days. **p**. Average investigation zone bout time across FR1 days. **b**,**g**,**h**,**m**,**o**,**p**. Grouped data are presented as means ± SEM. **c**,**d**,**f**,**i**,**j**,**k**,**l**. Data presented as boxplots with thick horizontal lines as respective group medians, boxes 25^th^ – 75th percentiles, and whiskers 1.5 x IQR. Individual data points as open circles.

We next examined social orienting in the presence of OTR blockade. We found that OTRA infusion did not influence interaction time (**Fig. 5J**), time spent attempting interactions (**Fig. 5K**), or total time in the interaction zone averaged across FR1 training (**Fig. 5L**). However, OTRA-infused mice traveled shorter total distances within the operant chambers on average across days, suggesting less exploration of the apparatus (**Fig. 5M,N**). This difference in activity was also reflected in fewer entries into the investigation zone by OTRA-infused mice compared to controls (**Fig. 5O**). This reduction in entries, however, was not substantial enough to alter the average amount of time the OTRA-infused mice spent in the zone per entry (**Fig. 5P**). Thus, OTR antagonism did not influence social orienting in this task, but may have reduced overall activity.

Next, we assessed how blockade of OTR influences an animal’s willingness to increase effort for access to a social partner. The PR3 testing revealed a lower breakpoint in the OTRA-infused mice compared to vehicle-infused controls (**Fig. 6A**), an effect which was driven by the females (**Fig. 6B**). There was no difference in the % successful rewards between groups (**Fig. 6C**), but OTRA-infused mice did show a reduction in interaction time (**Fig. 6D**), which was heavily influenced by their fewer rewards overall and mirrored their lower breakpoint (**Fig. 6E**). Despite this reduction in rewards and breakpoint, blocking OTRs did not influence the total time spent in the interaction zone (**Fig. 6F**), similar to FR1 performance. Unlike in FR1 training, the OTRA-infused mice did not differ in their activity levels as reflected by comparable distances traveled by the groups during PR3 testing (**Fig. 6G**). Together, these data indicate that blocking OTRs reduces, but does not completely abolish, social motivation by influencing reward seeking and not social orienting, especially in females.

**Figure 6.**
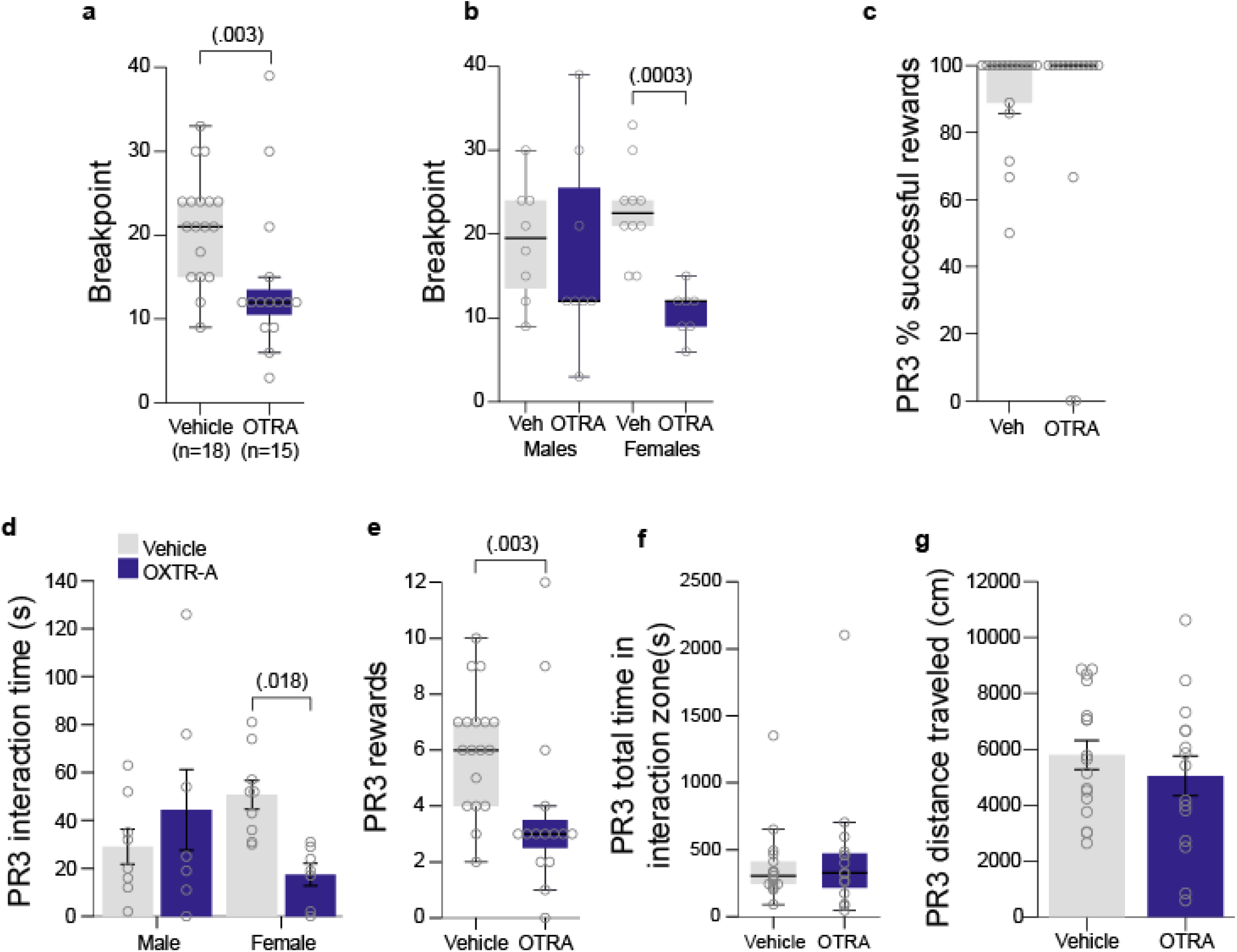
OTR antagonism reduces the motivation to achieve social interaction rewards. **a-b**. The breakpoint achieved during PR3 testing across infusion groups and segregated by sex. **c**. Percentage of successful rewards for both drug groups. **d**. Interaction time during PR3 testing. **e**. Number of rewards received during PR3 testing. **f**. Total time in the interaction zone during PR3 testing. **g**. Total distance traveled in the testing chambers by the OTRA and vehicle mice during PR3 testing. **d**,**g**. Grouped data are presented as means ± SEM. **a**,**b**,**c**,**e**,**f**. Data presented as boxplots with thick horizontal lines as respective group medians, boxes 25th – 75th percentiles, and whiskers 1.5 x IQR. Individual data points as open circles.

## Discussion

Here, we described in depth our novel social motivation assay that leverages operant conditioning to assess social reward seeking and social orienting, thus establishing a comprehensive assessment of social motivation behavior. We established in a commonly-used wildtype strain that mice will work via nosepokes for access to a transient social interaction reward and will work harder for a social interaction than for a non-social control reward. Further, we demonstrated a sex difference in this task, with male mice showing more robust social reward seeking and social orienting behavior compared to females. In addition, we provided two test cases for this assay: the *Shank3B* mutant with established sociability reductions and blockade of the prosocial oxytocin system.

Our task extends previous assays by enabling the dissociation of different aspects of social motivation as well as automating and simplifying the procedure. The addition of simultaneous video tracking allows for the quantification of movement and interactions with the stimulus animal, thus enabling interpretation of nosepoking behavior in light of social engagement - a limitation acknowledged in previous studies (Martin et al., 2014). Quantification of social interactions also allows us to disentangle the reward seeking aspects of social motivation from social orienting. This will enable future studies further dissecting the neural circuitry of these different social motivation components and investigating how genetic liabilities may differentially influence them. Extending the video tracking beyond previous work (Hu et al., 2021) to include that of the stimulus mouse enabled us to quantify additional metrics such as the duration and frequency of interactions between the stimulus and test animals, percentage of rewards with a successful interaction, and time spent attempting an interaction independent of the stimulus mouse behavior. This will allow for flexibility in future study designs, permitting differences in stimulus mouse behavioral responses to be explored. In addition, we leveraged the mouse’s innate tendency to nosepoke in dark holes, a more naturalistic behavior than lever-pressing, which allowed us to use the shorter 10 day protocol. Thus, our task more easily fits into an extensive ASC-model phenotyping pipeline. Further, a shorter protocol allows for social motivation assessment at earlier ages, such as during the social reward critical period (Nardou et al., 2019), whereas a full month of training would age an animal to early adulthood by the time of progressive ratio testing.

It should be noted, however, that a limitation of this task is the requirement of intact operant conditioning capabilities. As noted by Martin & Iceberg (2015), this task requires that the mice learn an association between the nosepoking behavior and the social reward. Thus, interpretation of the task will be limited in models with impaired conditioning abilities. However, coupling this task with another non-social operant conditioning paradigm (e.g. for food rewards) would provide a control for general conditioning ability and clarification as to whether any deficits are global or specific to the social domain.

Sociability deficits in *Shank3B* mice have been well-characterized (Guo et al., 2019; Peça et al., 2011; Wang et al., 2017). *Shank3B* KO mice display impairments in free socialization, approach of a social partner, and recognition of social novelty, especially in male animals. To assess sociability in our *Shank3B* mutants, we utilized video tracking during the habituation period. We found that, as in the literature, KO mice approached the social stimulus less than WT, and KO males spent less time in proximity to the social stimulus door than either WT or Het littermates. Thus, general sociability is indeed reduced in the KOs.

When tasked with learning to nosepoke for social reward, we found that all *Shank3B* genotypes could distinguish between the active and inactive holes, but KO mice nosepoke less, attain fewer rewards, and show less exploration of the apparatus. Overall, social reward appears to be less motivating to KO mutants than their Het and WT littermates. Previous work in another SHANK3 deficiency model found that *Shank3* mutants were less incentivized to overcome aversive environments for social reward (Rein et al., 2020). While this task was designed to measure social preference rather than motivation, their work aligns with our findings. Unfortunately, their study utilized only male mice, so it is uncertain whether the task would reveal similar sex differences as we observed in the current study.

Interestingly, here we observe a previously unreported phenotype in the Het mutants, which lack only one functional copy of *Shank3B* and thereby better model haploinsufficient patient populations. While we expected that fewer KOs would reach conditioning criteria, we were surprised that fewer Hets reached criteria as well. Further, as in the KO males, Het males also exhibited reduced social orienting. Moreover, Het males and KO males both had significantly lower breakpoints than WT males. This suggests that our assay has high sensitivity for detecting social motivation deficits in ASC-associated mouse models and represents a valuable addition to existing assays of social behavior. We anticipate that, particularly when measures of social orienting are included, studies of social motivation may broaden the field’s understanding of even well-characterized mouse models.

Disruption of oxytocinergic activity blunted wildtype animals’ motivation to work for a social interaction, including reward seeking behavior, yet did not influence social orienting as the mice spent comparable time near the social stimulus door. This is in line with previous work showing intact social approach behavior in oxytocin null mice, indicating this peptide is not essential for general spontaneous social approach in mice (Crawley et al., 2007). Increasing salience of, and thus attention to, social information via altered inhibitory tone has been suggested as a secondary, neuromodulatory role for oxytocin (Froemke and Young, 2021). Therefore, instead of blunting any social orienting behavior, it is more likely blockade of OTRs during our social motivation operant assay resulted in a reduction in salience of the social reward thereby decreasing motivation to achieve social interactions, reflected in fewer rewards. In the mesolimbic reward system, oxytocin has been shown to influence midbrain dopamine activity through modulation of GABAergic neuronal firing. Specifically, oxytocin biases activity towards the ventral tegmental area (VTA) and away from the substantia nigra, thus preferentially activating the reward system (Xiao et al., 2017). It is possible that preventing this shift in activity results in the reduced reward seeking behavior observed in OTRA mice. Yet, the OTRA mice did still show some social reward mediated behaviors. This could either be due to additional non-OXT circuits, or the fraction of OXT-responsive VTA dopamine neurons that are insensitive to OTR blockade, likely due to cross-reactivity with the vasopressin receptor (Xiao et al., 2017). Future work employing circuity mapping tools in OXT neurons will further clarify the role of oxytocin in social motivation.

Another opportunity apparent from our data arises from the fascinating propensity of unmanipulated, wildtype C57BL/6J littermates to dichotomize into two groups: socially motivated learners and socially unmotivated non-learners. Other behavioral paradigms similarly find such apparently inherent dichotomization of littermates into distinct behavioral phenotypes (Ellenbroek et al., 2005; Krishnan et al., 2007; Nestler and Waxman, 2020). Here, we have further repeated the PR3 test over multiple days which showed a remarkably stable intra-individual correlation (>.82), indicating that social motivation, at least in this task, exhibits a trait-like stability. This stability highlights an opportunity for investigations comparing highly motivated learners and their unmotivated non-learner littermates to further dissect the molecular underpinnings of social motivation.

Finally, though primarily we had selected our manipulations of OXT blockade and *Shank3b* mutation as test-cases for this new approach, synthesizing across all three studies included here does lead us to suggest one more novel conclusion: the circuitry that mediates social motivation may be sexually dimorphic in the mouse brain. Not only do WT males show greater levels of social orienting (e.g., interaction time), and reward seeking (e.g, total nosepokes in PR3), our two manipulations have disparate consequences on the sexes. Specifically, the female mouse social motivation circuit appears to be more dependent on OXT signaling in our assay, while only the male mouse social motivation circuit was vulnerable to *Shank3B* mutation. Such a ‘double dissociation’ is a classic demonstration that the circuits in the two sexes are distinct biological systems. Furthermore, of the two ASC-associated mouse models we have evaluated so far - *Shank3B* presented here and the previously-tested ASC/intellectual disability associated gene *Myt1l* (Chen et al., 2021) - both showed social motivation deficits only in male mice, while overall sociability measured by social approach was reduced in both males and females. This suggests this male specific vulnerability (or female protection) may be present across multiple models of ASC associated gene mutation. This sex difference in circuits is intruiging given the increase risk of ASC in males and could be further leveraged to understand the neurobiology of sex difference in social motivation, especially if this female protection replicates across many more models of the condition.

Overall, the protocol presented here expands existing assays by coupling automated, quantitative measures of social reward seeking with simultaneous measurement of social orienting. Social operant conditioning protocols have already proven useful in probing at inherent social differences between mouse strains and uncovering novel circuits underlying social motivation (Martin et al, 2014; Hu et al, 2021). We anticipate that our assay will provide further avenues for investigations of social motivation circuitry and the impact of ACS-associated gene mutations on social motivation. Future studies evaluating the rewarding properties of both social and non-social stimuli across multiple models will enable us to explore whether reduced social motivation is universally present or limited to a subset of ASC genetic liabilities, and to investigate whether deficits in reward processing are global or specific to the social domain. Finally, using the approach as part of the toolkit to examine the interaction between ASC risk factor models and sex might provide insights into the social circuits that are uniquely vulnerable in the male brain.

## Methods

All experimental protocols were approved by and performed in accordance with the relevant guidelines and regulations of the Institutional Animal Care and Use Committee of Washington University in St. Louis and were in compliance with US National Research Council’s Guide for the Care and Use of Laboratory Animals, the US Public Health Service’s Policy on Humane Care and Use of Laboratory Animals, and Guide for the Care and Use of Laboratory Animals.

### Social Operant Conditioning Task

Social motivation was evaluated in mice using a social operant conditioning task adapted and extended from previous methods (Martin et al., 2014; Martin and Iceberg, 2015) by adding continuous video tracking to measure, in parallel, both social reward seeking and social orienting aspects of social motivation (Chevallier et al., 2012). Each standard operant conditioning apparatus was enclosed in a sound-attenuating chamber (Med Associates) and modified to include a 3D printed filament door attached via fishing wire to a motor (Longruner), allowing it to be raised and lowered. This door was controlled by an Arduino (UNO R3 Board ATmega328P) programmed with custom code, connected to the Med Associates input panel. A clear acrylic conspecific stimulus chamber (10.2 × 10.2 × 18.4 cm; Amac box, The Container Store) was attached to the side of the operant chamber, centered between the nosepoke holes and separated from the chamber itself by a doorway (10.2 × 6 cm) containing vertical stainless steel bars spaced 6mm apart (**Fig. 1B**). The chamber was illuminated with white light during testing and an additional red chamber light illuminated in the apparatus during testing. The bottom tray of the operant apparatus was filled with one cup of fresh corn cob bedding, which was replaced between mice. Each operant chamber and stimulus chamber were designated for either males or females throughout the experiment. The operant chambers were cleaned with 70% ethanol and the stimulus chambers were cleaned with 0.02% chlorhexidine diacetate solution (Nolvasan, Zoetis) between animals.

Each operant chamber was outfitted with two nosepoke response holes. One was designated the active hole, which triggered illumination of a cue light within that hole and the raising (opening) of the door between the operant and stimulus chambers. The other hole was inactive and did not trigger any events. Designation of active and inactive status to right or left holes was randomized and counterbalanced across groups. When the door opened following a nosepoke in the active hole, the experimental and stimulus animals were able to interact through the bars for a 12-second social interaction reward before the door lowered (shut) and the active hole cue light turned off. The operant apparatuses were connected to a PC computer via a PCI interface (Med Associates). MED PC-V software quantified nosepokes as “active”, “inactive”, and “reward” based on the location of the poke and whether it occurred within an ongoing 12-second reward period (i.e. active nosepokes made while the door was raised were added to the “active” count but not the “reward” count). CCTV cameras (Vanxse) were mounted above the chambers and connected to a PC computer via BNC cables and quad processors. Ethovision XT v15 software (Noldus) was used to track the experimental and stimulus animals’ movement to quantify distance traveled, as well as time spent in and entries into the social interaction zones (defined as 6 × 8 cm rectangles in front of the door within both the operant and stimulus chambers). Custom Java and SPSS scripts were used to align the Ethovision tracking data with the timing of rewards in the Med Associates data to determine presence or absence of each animal within the interaction zones during each second of every reward period.

The operant conditioning protocol included phases of habituation, fixed-ratio conditioning (“training”), and progressive-ratio conditioning (“testing”) (**Figs. 1A, 3A, 4A**). For all trials, sex- and age-matched novel C57BL/6J mice served as conspecific stimulus animals. The stimulus mice were loaded into and removed from the stimulus chambers prior to the placement and after removal of the experimental mice into the operant chambers, respectively, to prevent the experimental animals from being in the chambers without a conspecific stimulus animal present. Habituation consisted of 30 minute sessions on each of two days, during which the door remained open and the nosepoke holes were inaccessible. This allowed the experimental mice to acclimate to the chamber and to the presence of a stimulus partner in the adjoining chamber, as well as providing an opportunity for a measure of social approach behavior. Fixed- and progressive-ratio conditioning consisted of 1-hr daily sessions during which nosepokes in the active hole were rewarded with a 12-second social interaction opportunity. During the 12-sec reward period, any additional active nosepokes did not result in another reward.

Several schedules of reinforcement were utilized across the conditioning phases. A fixed-ratio schedule of one reward per one active nosepoke (FR1) was employed for at least the first three days of training, after which mice were individually advanced to the next phase upon achievement of conditioning criteria. Conditioning criteria were defined as, within a single session, achieving at least 40 active nosepokes, a 75% poke accuracy, and 65% successful rewards (defined as both experimental and stimulus mice in their respective social interaction zones simultaneously for at least 1 sec of the reward, or time at the door for the control group in cohort 1). Animals who failed to meet these criteria after 10 days (based on cohort) of FR1 were designated as non-learners. For animals that successfully met conditioning criteria, FR1 was followed by three days of a fixed ratio 3 (FR3) training, in which three active nosepokes were required to obtain a single reward. All mice then received 1 to 4 days of a progressive ratio 3 (PR3) testing to determine their breakpoint, or maximum effort the animal will exert for a social interaction. In this phase, rewards became progressively more effortful to obtain, with three additional nosepokes required for each subsequent reward (i.e. 3, 6, 9, 12, etc). Breakpoint was defined as the number of rewards after which the animal would no longer continue to engage in nosepoking behavior.

### Animals

All mice used in this study were maintained and bred in the vivarium at Washington University in St. Louis. For all experiments, adequate measures were taken to minimize any pain or discomfort. The colony room lighting was on a 12:12h light/dark cycle; room temperature (∼20-22°C) and relative humidity (50%) controlled automatically. Standard lab diet and water were freely available. Upon weaning at postnatal day (P)21, mice for behavioral testing were group housed according to sex and experimental condition.

Social motivation was assessed in three separate cohorts. The first cohort included unmanipulated wildtype C57BL/6J mice (20 females, 20 males) that served to validate the task. Twenty mice (10 females, 10 males) served as experimental mice that received transient access to a social partner as a reward during testing. The remaining 20 mice (10 females, 10 males) served as control mice that received as a reward only the opening of the door to expose a metal apparatus wall. The testing procedure for this cohort was as stated above, except that all mice received up to 18 FR1 training sessions and 4 PR3 testing sessions to assess reliability of performance within individuals (**Fig. 1A**). All mice were young adults (*M*=P68, range P56-P82) at the start of testing.

The second cohort consisted of mice modeling genetic liability for Phelan-McDermid Syndrome, a neurodevelopmental disorder characterized by global developmental, speech and motor delay, intellectual disability, and high prevalence of ASC (Kolevzon et al., 2014). Specifically, these C57BL/6J mice harbor a disruption to the PDZ domain of the *Shank3B* gene (Peça et al., 2011). To generate this cohort, we crossed *Shank3B* heterozygous mutants (*Shank3B* Het) producing 14 wildtype (*Shank3B* WT; 7 females, 7 males), 10 *Shank3B* Het (5 females, 5 males), and 14 *Shank3B* homozygous mutant (*Shank3B* KO; 6 females, 8 males) littermates. The testing procedure for this cohort was as stated above, with 10 days of FR1 testing since more than 85% of experimental mice in the first cohort reached conditioning criteria by then (**Fig. 3A**). All mice were young adults (*M*=P79, range P70-P99) at the start of testing.

The third cohort comprised 33 C57BL/6J wildtype mice administered either an oxytocin receptor antagonist (OTRA; n=15; 7 females, 8 males) or vehicle-only (n=18; 10 females, 8 males) on each day of the protocol (**Fig. 5A**). Drug administration details are described below. The testing procedure for this cohort was as stated above, with 12 days of FR1 testing since more than 85% of experimental mice in Cohort 1 reached criteria before then. All mice were young adults (P69-P72) at the start of testing.

### Intracerebroventricular infusion of oxytocin receptor antagonist

Surgeries were conducted at P59. Twenty-four hours prior to surgery, each animal was given 0.25 mg of the chewable anti-inflammatory Rimadyl (Bio-Serv, Flemington, NJ) and the surgical area was shaved. Mice were anesthetized with 2.5-5% isoflurane and placed in a stereotaxic apparatus. Mice received a local anesthetic, 1 mg/kg of Buprenorphine SR (ZooPharm, Laramie, WY), an antibiotic, 2.5-5 mg/kg of Baytril (Bayer Healthcare LLC, Shawnee Mission, KS), and 0.5mL of 0.09% sterile saline. Following cleaning of surgical area by alternating 70% Ethanol and Betadine surgical scrub (Purdue Products L.P., Stamford, CT) three times, an incision was made along the skull and skin retracted to visualize bregma to lambda. The periosteum was removed by lightly scratching the surface of the skull and the area was cleaned three times with a betadine solution, followed by 3% hydrogen peroxide. The guide cannula was cut to a length of 2mm, so that it would enter the lateral ventricle, and placed in a stereotaxic cannula holder (Stoelting, #51636-1). Using a rapid, fluid motion, the 26-gauge unilateral guide cannula (C315GS-5/SPC, Plastics One, Roanoke, VA) was inserted at the following coordinates: M/L=+1, A/P=−0.4, D/V −2.2, based on prior work (Mantella et al., 2003; Radulovic et al., 1999; Raposinho et al., 2001). C&B Metabond dental cement (Parkell, Edgewood, NY) was mixed on a chilled ceramic dish and used to secure the cannula to the skull and seal the surgical area. The dental cement dried completely and a dummy cap was inserted to prevent clogs of the implanted cannula. The dummy cap (C315DCS-5/SPC) and internal cannula (C315IS-5/SPC) were cut to protrude 0.2mm from the end of the guide. The animal was removed from the stereotaxic apparatus and placed in a recovery cage. Animals were housed together after fully awake and provided an additional 0.25mg dose of Rimadyl. During daily monitoring, dummy caps were replaced and tightened as needed. Mice were given seven days to recover prior to testing, and were euthanized at any sign of distress or damage to the surgical area.

All mice received 4µl infusions at least one hour before participating in the social operant task each day. Mice were randomly assigned to receive either vehicle (artificial cerebrospinal fluid solution, Tocris Bioscience, Bristol, UK; n=18) or an oxytocin receptor antagonist (OTA; desGly-NH_2_,d(CH_2_)_5_[Tyr(Me)^2^Thr^4^]OVT, Bachem, Torrance, CA; n=15). The OTA, dissolved in vehicle at 0.25 ng/uL, is a peptidergic ornithine vasotocin analog chosen because of its broad applicability and prior use in ICV injections (Ferguson et al., 2001, 2000). The solutions were delivered into the lateral ventricles through a 33-gage internal cannula (C315IS-5-SPC) via a PlasticsOne Cannula Connector (C313CS) over the course of 120 seconds using a programmable syringe pump (New Era pump systems #NE-1200). Mice were restrained by neck scruff while the internal cannula was inserted and then were placed in a clean holding cage, allowing free movement during infusion. To ensure complete diffusion, the internal cannula remained inserted for 60-90 seconds post infusion, after which mice were returned to home cage. Following completion of all behavioral testing, cannula placement was confirmed by infusing 10ul of dye (India Ink, Higgins, Leeds, MA) to flood the ventricles and immediately euthanizing the animal via isoflurane overdose. Brains were extracted and sliced coronally at the injection site with a razor blade. Infusion of the dye into the ventricles was then confirmed by eye and infusions that missed the ventricles were excluded from the final analysis.

### Statistical Analysis

Statistical analyses and data visualization were conducted using IBM SPSS Statistics (v.27). Prior to analyses, data were screened for missing values and fit of distributions with assumptions underlying univariate analysis. This included the Shapiro-Wilk test on z-score-transformed data and qqplot investigations for normality, Levene’s test for homogeneity of variance, and boxplot and z-score (±3.29) investigation for identification of influential outliers. Means and standard errors were computed for each measure. Analysis of variance (ANOVA), including repeated measures, was used to analyze data where appropriate, and simple main effects were used to dissect significant interactions. Sex was included as a biological variable in all analyses across all experiments. Where appropriate, the Greenhouse-Geisser or Huynh-Feldt adjustment was used to protect against violations of sphericity. Multiple pairwise comparisons were subjected to Bonferroni correction, where appropriate. For data that did not fit univariate assumptions, non-parametric tests were used or transformations were applied. Sex*genotype effects are reported where significant, otherwise data are reported and visualized collapsed for sex. The critical alpha value for all analyses was p < .05 unless otherwise stated. Figure illustrations were generated using BioRender. The datasets generated and analyzed during the current study are available from the corresponding author upon reasonable request. Details of all statistical tests and results can be found in Tables 2-4.

## Acknowledgements

This work was generously supported by the NICHD P50HD103525 (IDDRC@WUSTL), NIMH R01MH107515-05 and R01MH124808 (JDD). We would like to thank Dr. John Constantino for guidance, Drs. Adrian Gomez, Kyle Parker, Ream Al-Hasani, and Daniel Castro for operant paradigm training, Kyle Kneipkamp for assistance designing and creating custom chambers, and Dr. Elena Minakova and Kayla Nygaard for surgery training.

## Statistical Tables and Supplemental Figures

**Figure S1.**
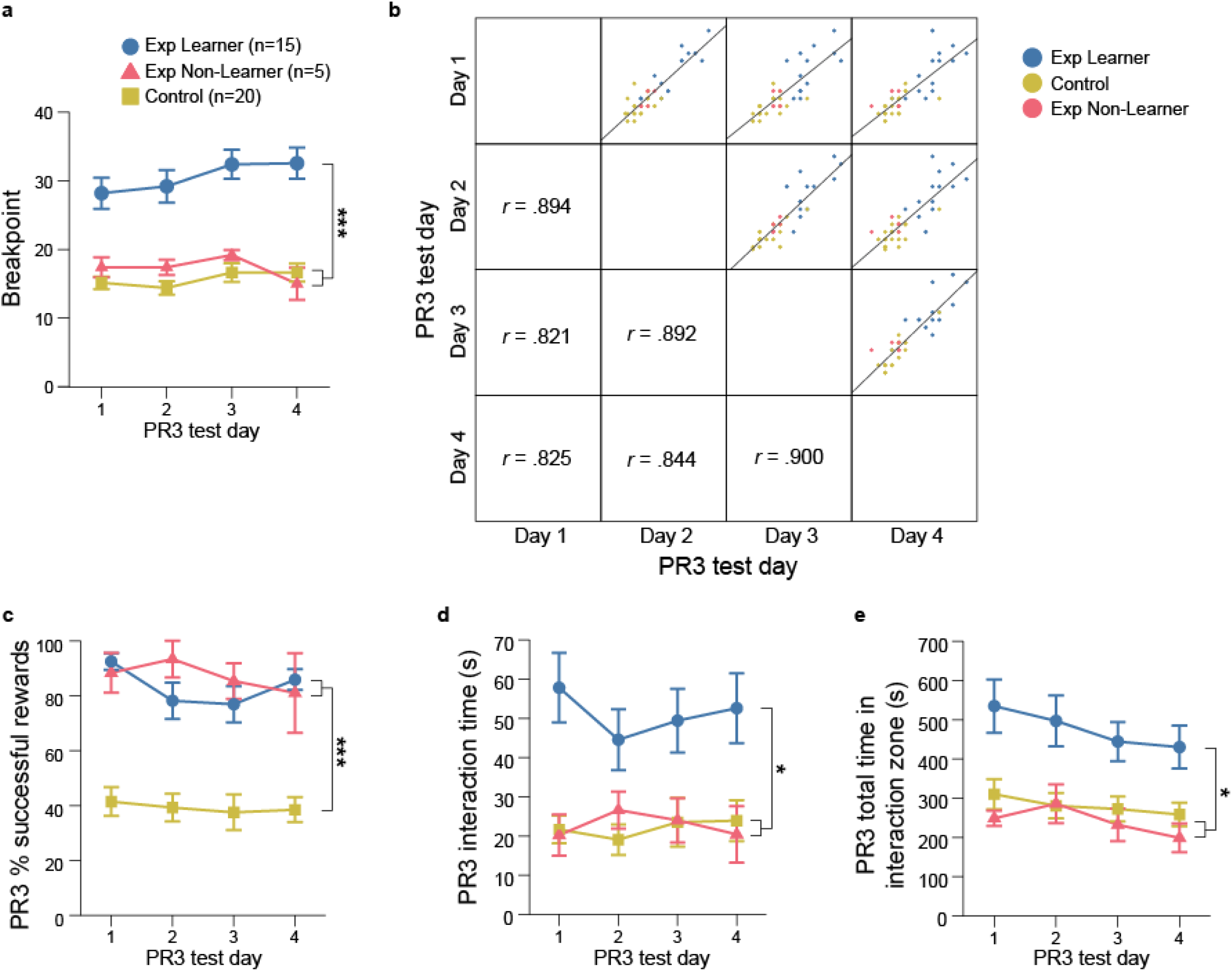
Social motivation levels are highly stable within an individual mouse. **a**. Breakpoints achieved across each day of PR3 for experimental learners (Exp Learners; n=15), experimental non-learners (Exp NL; n=5) and control (Control; n=20) non-learners. **b**. Correlation matrices between breakpoints achieved on all four days of PR3 testing. **c**. The percentage of successful rewards for each group across each day of PR3. **d**. Interaction time across all days of PR3 for all groups. **e**. Total time in the interaction zone across all days of PR3 for all groups. **a**,**c-e**. Grouped data are presented as means ± SEM. Statistical significance, ***p<.001, **p<.01, *p<.05.

**Figure S2.**
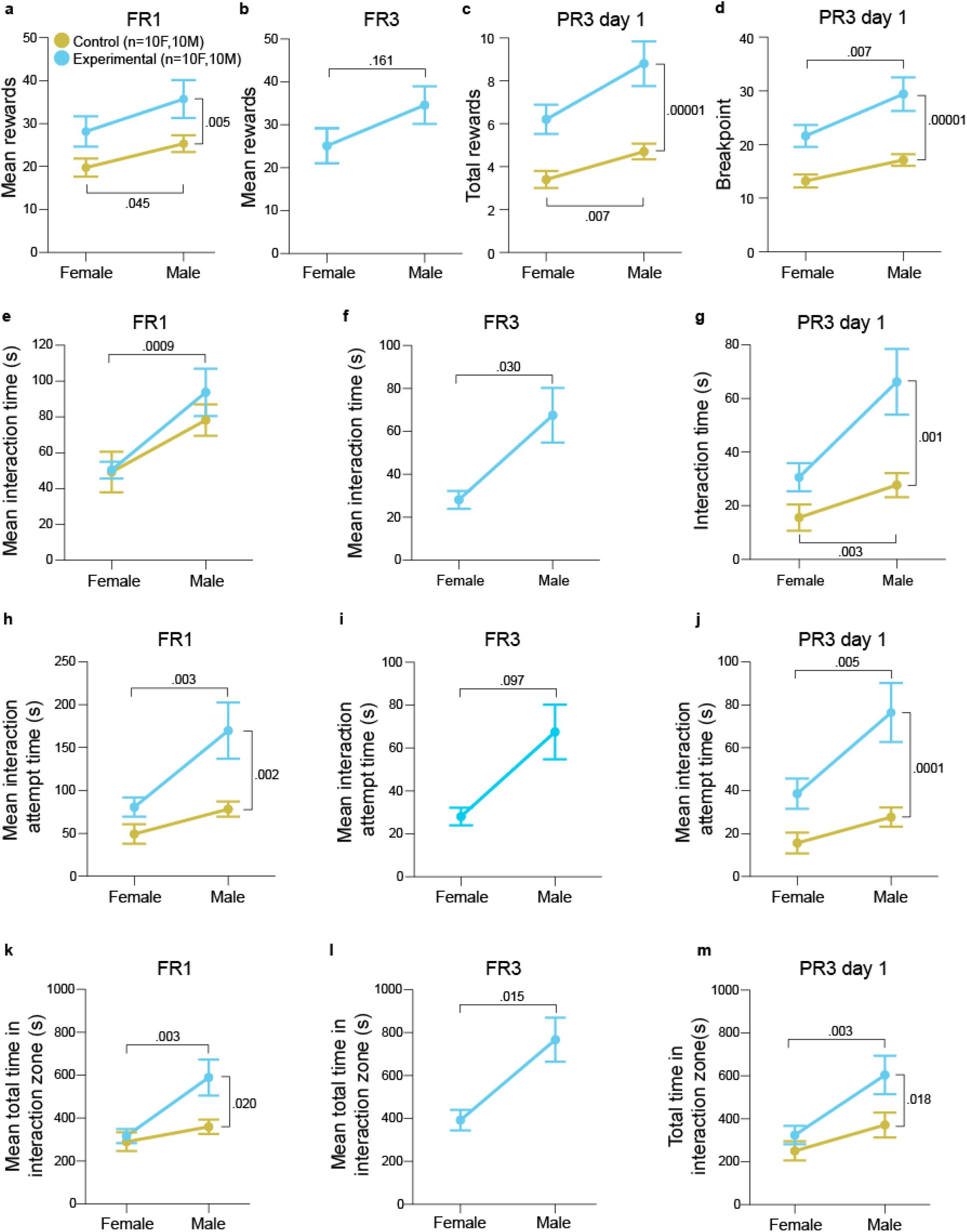
Male mice exhibited increased reward seeking and social orienting. **a**. Number of average rewards received across FR1 training by experimental and control male (n=20) and female (n=20) mice. **b**. Average number of rewards received across all three FR3 days by experimental and control male and female mice. **c**. Total rewards received by male and female mice of each group during PR3 day 1. **d**. Breakpoints achieved by male and female mice of each group on day 1 of PR3 testing. **e-g**. Interaction time averaged across **(e)** FR1 days, **(f)** FR3 days, and **(g)** for PR3 day 1 in female and male mice of both groups. **h-j**. Interaction attempt time averaged across **(h)** FR1 days, **(i)** FR3 days, and **(j)** PR3 day 1 for female and male mice of both groups. **k-m**. Total time in the interaction zone averaged across **(k)** FR1 days, **(l)** FR3 days, and **(m)** PR3 day 1 for female and male mice of both groups. For all panels, the grouped data are presented as means ± SEM. Statistical significance, ***p<.001, **p<.01, *p<.05.

**Figure S3.**
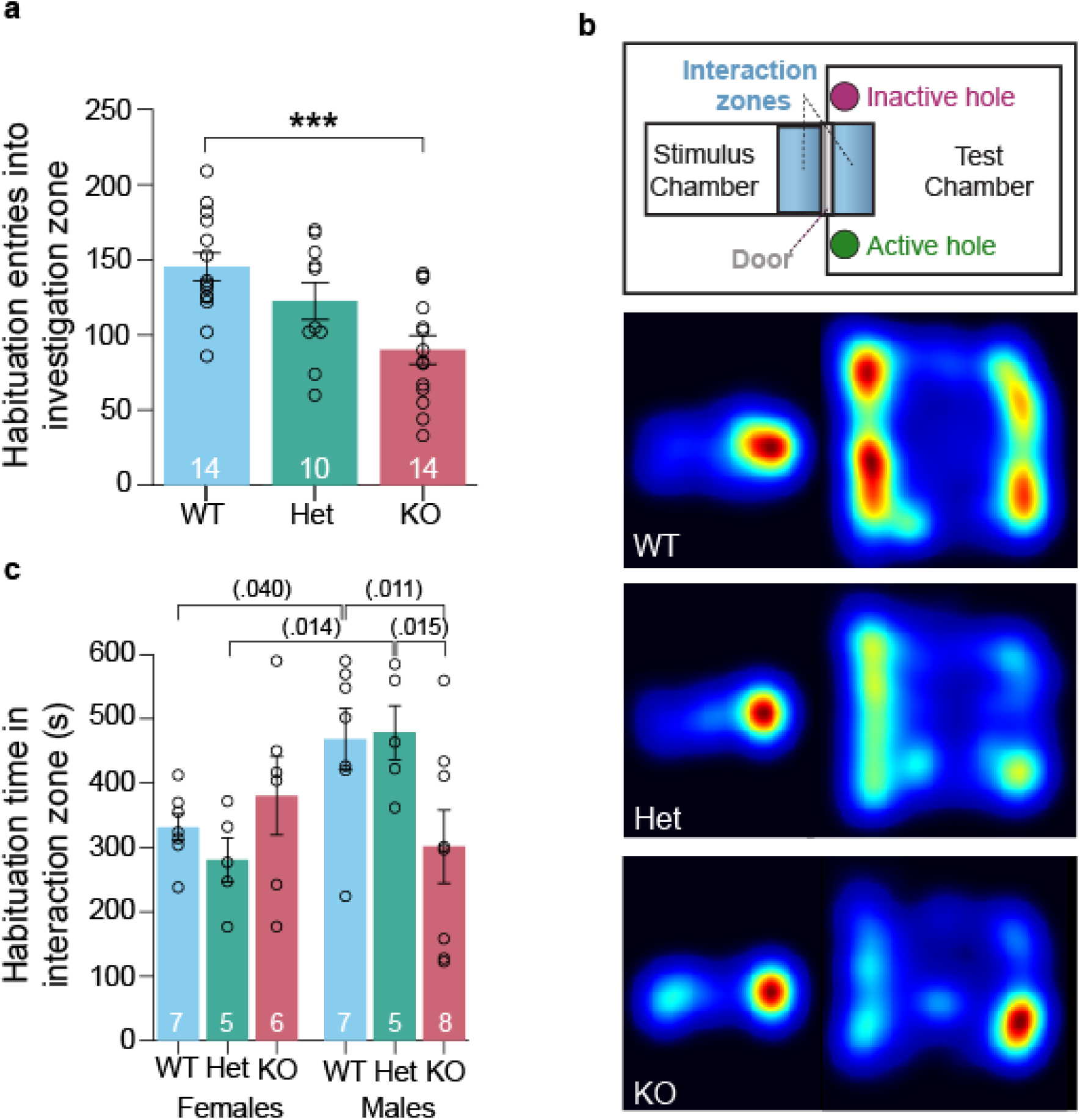
*Shank3B* knockout (KO) mice exhibit reduced social approach during the habituation period relative to wildtype (WT) and heterozygous (Het) mice. **a**. Number of entries into the interaction zone during day 2 of habituation for WT (n=14), *Shank3B* Hets (n=10), and KO mice (n=14). **b**. Schematic of apparatus (top panel) and representative heatmap of the animal’s position in the apparatus during habituation for each genotype. Warmer colors represent greater time spent in that position. **c**. Time in the interaction zone during day 2 of habituation habituation split by sex (WT, females = 7; Het, females = 5; KO, females = 8). **a-b**. Grouped data are presented as means ± SEM with individual data points as open circles. Statistical significance, ***p<.001, **p<.01, *p<.05.

